# Lacuna: Cryptic Binding Pocket Discovery via Conformational Ensemble Analysis

**DOI:** 10.64898/2026.08.14.744956

**Authors:** Clayton W. Moore

## Abstract

Lacuna, an open-source Python tool for discovering cryptic binding pockets: sites that are absent or too small to detect in the unbound structure of a protein, RNA or DNA and open only during conformational fluctuation. Most binding-site predictors score a single static structure, the very structure in which a cryptic site is invisible. Lacuna generates a conformational ensemble from any input structure, detects pockets independently in every conformer, clusters the detections into persistent sites, and ranks those sites with a model fitted on within-structure pairs. Ensemble generation is pluggable: normal mode analysis (NMA) by default, with implicit-solvent molecular dynamics, Boltz-2 diffusion sampling or a user-supplied ensemble as alternatives. On the designated test fold of CryptoBench, all 222 structures of which are scored, the zero-dependency default recovers the cryptic site for 59.9% of structures in its top five predictions, and it recovers 69% on the PocketMiner set, 41% on a curated set of literature apo/holo pairs and 89% of structures in the already-open COACH420 set. On that test fold, ordering the candidate set with the fitted ranker rather than the analytic rule it replaced accounts for a gain of 27.5 points (95% CI +21.2 to +33.8), the largest single effect measured here. An optional protein-language-model (PLM) ranker reaches 66.1% on a 180-structure subset of that fold, and pooling the geometric detector with an optional learned surface detector raises top-five recovery to 73.9% and the fraction of structures in which the site is found at all to 86.4% in a separate held-out evaluation at five conformers (184 structures; both optional configurations measured with version 1.4.0). A new per-residue cryptic-site propensity reaches AUROC 0.766 and AUPRC 0.340 on the same test fold (231 targets), against 0.249 for the same scorer on the input structure alone, so the ensemble carries information the input does not. RNA and DNA need no extra flags; on RNA sites with no cavity in the input the ensemble raises the fraction found from 44% to 73% (measured with version 1.3.0). With the default NMA backend the pocket report is identical at any thread count and between Windows and Linux on the 40 proteins compared, ensemble generation needs less than half the memory of version 1.4.0 from 1,300 residues up, and the full pipeline is 7% to 30% faster than 1.4.0 on the reference proteins of the repository’s speed table. At the default 20 conformers it takes a median of 1.9 seconds per chain on one CPU thread (30 chains of 57 to 676 residues). Outputs include docking-ready Boltz constraints and AutoDock Vina boxes, a per-residue propensity map and the conformer ensemble. Lacuna is MIT licensed, available at https://github.com/mooreneural/lacuna and on PyPI as lacuna-pockets, and runs as a local MCP server for AI assistants and on hosted no-install platforms.

## 1 Introduction

Many disease-relevant human proteins are classified as undruggable because their experimentally determined structures present no well-formed pocket for a small molecule to occupy [1]. A substantial fraction of those are not truly featureless: they carry cryptic sites, pockets that are closed in the apo structure and open on ligand binding or thermal fluctuation [2]. K-Ras was considered undruggable for three decades until a cryptic pocket beneath switch-II was identified, which led directly to a clinical programme [3]. The practical question is not whether such sites exist but whether they can be found computationally before a ligand is known.

This is hard for a specific and structural reason. The dominant approach to binding-site prediction scores a single static structure, whether geometrically by identifying concave surface features [4] or by machine learning over surface descriptors [5] or learned residue representations [6]. Applied to an apo structure containing a cryptic site, these methods are being asked to find a cavity that is not there. Methods built specifically for cryptic sites predict per-residue opening propensity from a single structure [7], which sidesteps the geometry problem but returns residue scores rather than a pocket a docking program can accept.

The alternative is to supply the missing conformations. Molecular dynamics followed by pocket detection on the trajectory is the established route [8], and it works, but it places a simulation budget between the user and an answer, which in practice restricts it to targets already considered worth the investment. That cost is why ensemble-based pocket finding has not become routine tooling in the way single-structure detectors have.

> **Key Problem.** A cryptic site is absent or poorly formed in the apo structure, so a detector given one static structure is asked to find a cavity that is not there. Molecular dynamics exposes the missing conformations but puts a simulation budget between the user and an answer, which keeps ensemble-based pocket finding off the default path for most targets.

Lacuna is built on the observation that the ensemble does not have to be expensive to be useful. A cryptic pocket needs only to open somewhere in a set of plausible conformations for a geometric detector to see it, and the cheapest useful source of such conformations, an elastic network model, costs seconds rather than CPU-days. What matters is then the bookkeeping: detecting pockets per conformer produces a large, redundant set of transient cavities that must be matched across conformers into persistent sites and ordered so the interesting one appears near the top. The same reasoning holds for structured RNA, where a ligand site can likewise be closed in the deposited structure, and Lacuna applies the identical pipeline to RNA and DNA.

Our contributions are as follows.

### Lacuna makes ensemble-based pocket discovery cheap enough to be routine

The default backend requires no simulation setup, no force field parameterisation and no GPU. On one CPU thread the full pipeline takes a median of 1.9 seconds per chain at the default 20 conformers, and its sampler needs less than half the memory of version 1.4.0 from 1,300 residues up.

### Ensemble generation is pluggable, including a co-folding backend

Normal mode analysis, implicit-solvent molecular dynamics, Boltz-2 diffusion sampling, and arbitrary user-supplied ensembles all present the same interface, so the sampling method is a parameter rather than a rewrite.

### Transient pockets are first-class objects

Detections are clustered across conformers into sites that carry their own statistics: how often the site is open, how much it opens relative to the input structure, and how far its centroid wanders. These ensemble-derived quantities are unavailable to any single-structure detector and are what the ranker relies on most.

### Ranking is fitted, not hand-tuned

A linear model over 23 features, trained on within-structure pairs so that it optimises ordering directly, recovers the site for 59.9% of CryptoBench test-fold structures, all 222 of them, against 32.4% for the analytic crypticity rule it replaced, roughly twice as many.

### Detection is a pluggable choice, and pooling detectors raises coverage

Beyond the default geometric detector, an optional learned surface detector scores the probe-accessible surface directly and so proposes sites too closed for geometry to register. Pooling the two with --detector surface-fusion and ranking the union with a matching fitted model raises top-five recovery to 73.9% and the fraction of structures in which the site is found at all from 68.5% to 86.4% on a separate held-out set of 184 structures, measured at five conformers with version 1.4.0.

### Protein, RNA and DNA go through one pipeline

Nucleic-acid input is recognised from its residues and sampled with an elastic network built on three beads per nucleotide [17]; detection, clustering and ranking are shared with protein.

### The same input gives the same report on every machine

With the default NMA backend the pocket report is identical at any linear-algebra thread count for protein, RNA, DNA and symmetric assemblies, and between Windows and Linux on the 40 proteins compared, with nothing to configure.

### Output is docking-ready, and per residue as well as per pocket

Each site is emitted as a Boltz YAML constraint, an AutoDock Vina box, and a pseudoatom PDB, so a predicted pocket can be handed directly to a docking or co-folding run, and a per-residue propensity map places Lacuna on the per-residue scale most cryptic-site predictors are evaluated on.

### Lacuna runs where people already work

Beyond the command line and Python API, a local Model Context Protocol (MCP) server lets an AI assistant run Lacuna on the user’s own machine, the repository imports as a Claude skill, and hosted servers run it with no installation at all.

Lacuna is MIT licensed, distributed on PyPI, archived on Zenodo [9], and carries a test suite of 371 tests. This report describes the design and validates the implementation. A separate study uses Lacuna’s per-candidate output, alongside three other detectors, to analyse how the field’s standard evaluation metric conflates detection with ranking; that analysis is reported elsewhere [10] and is not restated here.

### How to interpret these results

Lacuna is built for cryptic-site discovery, not general binding-site prediction. Against P2Rank [5], a strong general-purpose predictor, the two tools are close on both cryptic and already-open sites, with no difference resolved at these sample sizes (Section 3.3). What the default adds is that it runs in seconds on a CPU with no external neural-network checkpoint, no MSA and no GPU, and that its 23 fitted coefficients ship in the source. The contribution is a modular ensemble-analysis system with ensemble-derived site properties and docking-ready output, not a detector that dominates all alternatives.

**Figure 1:**
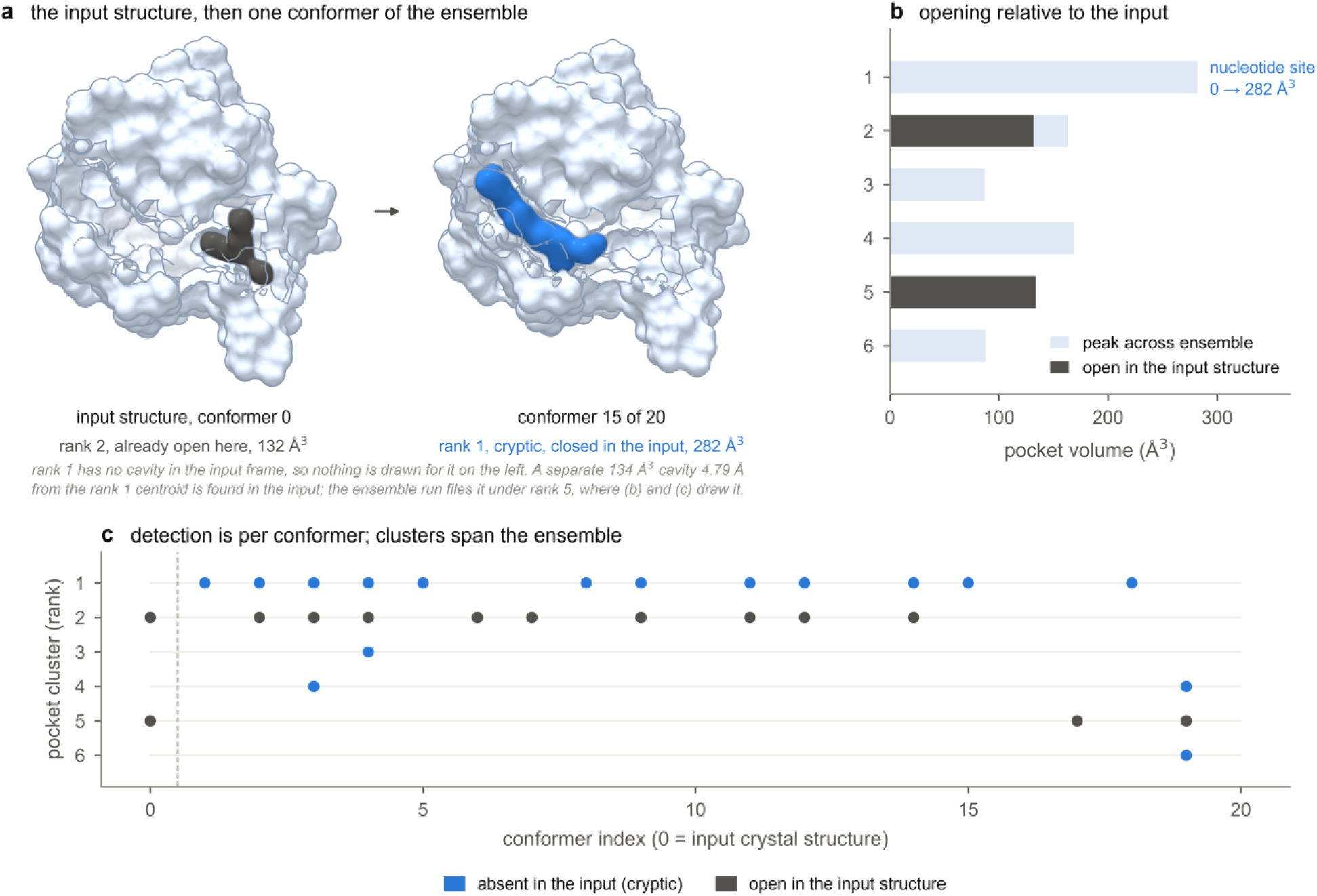
A site absent from the input structure, recovered at rank 1 in under a second. Lacuna run on chain A of apo K-Ras (PDB 4OBE) with default settings: normal mode backend, 20 conformers. (a) The structure itself, before and after. Left, the input crystal structure with the cavity that is already open there, the rank-2 cluster at 132 Å³, in dark grey. Right, conformer 15 with the rank-1 cryptic cavity at 282 Å³, in blue. Both frames are drawn on one camera and one crop, so the two are at one scale. One window serves the pair: it is the footprint of the cavities the pair draws, rank 2 in the input frame and rank 1 in conformer 15, and both frames are drawn under that same window, so the two differ only in the molecule. It leaves the molecular surface undrawn over the atoms that stand in front of it, 6.3% of atoms in the input frame and 7.4% in conformer 15, so that the cavity can be seen at all; that window is a drawing decision and not an opening in the molecule. Nothing is drawn where the detector found no cavity: it reports none for rank 1 in the input frame and none for rank 2 in conformer 15, and neither is drawn. Rank 1’s input volume of 0 Å³ is a statement about that cluster rather than about the input structure. A separate run on the input structure alone finds a 134 Å³ cavity whose centre is 4.79 Å from the rank-1 centroid, which the ensemble run files under rank 5: panel (b) plots it as the dark bar at rank 5 and panel (c) marks it at rank 5, conformer 0, and only the two renders omit it. (b) Pocket volume in the input structure against the peak volume reached anywhere in the ensemble. The rank-1 cluster goes from 0 to 282 Å³. Clusters 2 and 5 are ordinary surface pockets, already open in the input, and the ensemble adds little to them. The rank-1 cluster reaches a Jaccard overlap of 0.38 with the benchmark annotation for this target, which is residues 12, 13 and 25 to 36, and contains 79% of them. That annotation is the P-loop plus the switch I region, not the switch-II pocket. The switch-II pocket that sotorasib and adagrasib bind is the rank-2 cluster, which matches all seven of its lining residues (V9, Q61, R68, M72, H95, Y96, Q99) but already measures 132 Å³ in the input structure and so scores a crypticity of only 0.153. On this target Lacuna ranks an emptied ligand site first and the genuine cryptic pocket second. (c) Each row is a pocket cluster, each column a conformer, and a mark indicates the cluster was detected in that conformer. Column 0, left of the dashed line, is the input crystal structure itself; columns 1 to 20 are generated conformers. Clusters absent from the input structure are coloured blue. The rank-1 cluster is the nucleotide site: it is detected in 12 of the 20 generated conformers and in none of the input, which is what makes it look cryptic here and what makes it invisible to a detector that sees only column 0. Its lining is the P-loop (residues 10 to 18), the switch I stretch (28 to 38), which is eleven of its twenty-six lining residues and carries most of the overlap quoted above, the DxxG motif (57 to 60) and the first two residues of the NKxD motif (N116 and K117), and it reads as absent from the input because Lacuna discards HETATM records, so the bound GDP is removed and what the detector does find there in the crystal conformer, the 134 Å³ cavity named above, is clustered under rank 5 rather than with this site. Every value plotted and rendered here comes from one version 1.5.0 run, which takes 0.82 second of pipeline time on the desktop CPU described under Runtime and memory; the only quantities quoted from outside it are the 134 Å³ cavity, its 4.79 Å offset from the rank-1 centroid and its match to rank 5, which come from the separate zero-conformer run named above.

## 2 Design

### 2.1 Overview

Lacuna is a four-stage pipeline (Figure 2). A protein, RNA or DNA structure enters, an ensemble is generated, pockets are detected independently in each conformer, detections are clustered across conformers, and the resulting sites are ranked and written out. Each stage is separable, and the ensemble stage in particular is a plugin point. Protein, RNA and DNA share every stage, and with the NMA backend the report does not depend on thread count (Section 3.6).

#### Ensemble generation

The default backend is an anisotropic network model [11], an elastic network in which residues are nodes connected by harmonic springs. Displacing the structure along its lowest-frequency normal modes produces conformers that respect the fold’s intrinsic flexibility at negligible cost. This is deliberately the cheapest reasonable choice and also the most limited: harmonic modes describe collective breathing well and cannot produce large hinge motions or loop rearrangements. For RNA and DNA the network is built on three beads per nucleotide (C1’, C2 and P) at a 9 Å cutoff, the three-bead model of Pinamonti et al. [17]; it is selected automatically from the input’s residues, and --backend rna-nma requests it explicitly. Nucleic acids share the protein sampling loop, so everything downstream is identical.

The NMA backend is built so that its output does not depend on how the linear-algebra library is threaded. Every intermediate, from the Hessian through the eigendecomposition to the propagation of mode displacements to all atoms, is computed in double precision and cast to single precision once at the end; every mode is given a fixed sign; and modes that share an eigenvalue, as they do in any assembly with a three-fold or higher symmetry axis, are given a canonical basis computed from the space they span rather than whichever basis the solver returned. The Hessian is assembled a slab of rows at a time and factored in place, which is where the memory and time savings reported under Runtime and memory (Section 3.7) come from.

Three alternatives share the same interface. An implicit-solvent molecular dynamics backend samples thermal motion directly. A Boltz-2 backend [16] draws conformers as independent samples from a co-folding model’s learned posterior, with the diffusion step scale lowered from its default to increase structural diversity; its conformers are paired to the input by sequence order and superposed onto it before detection, and the backend remains experimental. It requests a multiple sequence alignment from the public ColabFold server by default, which --no-msa-server turns off. Finally, --ensemble accepts a multi-model PDB or a directory of structures, so an ensemble produced by any external method can be analysed without Lacuna generating anything. Imported conformers are superposed onto the input and paired with it by residue identity, falling back to residue order with a warning when a tool has renumbered them, so a conformer missing a loop remains usable. Because every quantity Lacuna reports is relative to the input, a sanity check warns when an imported ensemble no longer describes it, measuring drift, rigid-body offset, radius-of-gyration ratio and the spread between conformers.

#### Pocket detection

Within each conformer, pockets are found by a grid-based alpha-sphere method in the fpocket lineage [4]: candidate spheres are placed where they contact protein atoms without penetrating them, retained within a radius band that admits ligand-sized cavities while rejecting bulk solvent and interstitial voids, and clustered spatially into pockets. Lining residues are assigned by true atomic contact rather than by a radius around the pocket centre, which matters because a centre-and-radius definition inflates the residue set of large pockets and makes any overlap metric computed from it optimistic. Alternative detectors are available. A learned surface detector scores the probe-accessible molecular surface with a fitted model rather than requiring a formed concavity, so it can reach sites the geometric method discards as too shallow; --detector surface-fusion pools its proposals with the geometric detector’s and ranks the union with a model fitted on that combined candidate set. Its full configuration needs the same optional protein-language-model dependency as the PLM ranker, and a geometry-only variant (--no-sequence) runs without it at a smaller gain. --detector p2rank substitutes P2Rank [5] per conformer.

#### Cross-conformer clustering

This is the step that turns per-conformer detections into sites. Pockets from different conformers are matched into clusters by spatial proximity of their centres together with overlap of their lining residue sets, so that the same physical site detected in fifteen conformers becomes one cluster rather than fifteen candidates. Each cluster then carries statistics no single structure can provide: persistence, the fraction of conformers in which the site is open; the volume distribution across the ensemble and its coefficient of variation; the volume in the input structure specifically; and the positional spread of the cluster’s centroid between conformers. A site’s lining is the set of residues present in a strict majority of the conformers in which it is detected, which keeps linings close to the size of a real site: on 120 CryptoBench test targets the median lining is 18 residues and 1.27 times the size of the annotated site, with no loss of top-five recovery.

#### Ranking

The default strategy is a linear model over 23 features spanning pocket geometry, druggability in the sense of Halgren’s descriptors [12], and the ensemble-derived terms above. It is fitted on within-structure pairs, so it optimises the ordering of candidates inside one protein rather than classifying pockets in isolation, which is the quantity that actually determines whether a user sees the right site. Fitting used only CryptoBench’s homology-separated training folds. An optional strategy, learned-plm, refits the same linear form with four additional features summarising a protein language model’s per-residue cryptic-site probabilities over the cluster’s lining [13]; it requires an optional dependency and is not the default, so that the same command produces the same ranking on every installation. The model fitted for the pooled detector, which --detector surface-fusion selects automatically, can also be named directly with --rank-by learned-fused. Several analytic strategies remain available for targets that resemble classical case studies more than they resemble CryptoBench.

#### Crypticity

Independently of rank, every site receives a continuous score between 0 and 1 capturing the conformational-selection signature:

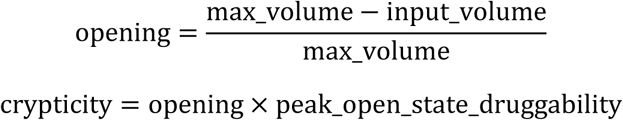

A site absent from the input structure has opening 1.0, so crypticity reduces to its druggability once open. A site already open in the input has low opening and therefore low crypticity regardless of how druggable it is. A site is flagged cryptic when its crypticity clears a fixed threshold of 0.3.

#### Outputs

Sites are written as a ranked JSON pocket report carrying every quantity above, together with each site’s chain composition and a flag for sites at a chain interface, and optionally as Boltz YAML constraint files, AutoDock Vina box configs, pseudoatom PDB files for visualisation, and the generated conformational ensemble as a multi-model PDB (--emit-conformers) for reuse in docking or molecular dynamics. The report also carries a per-residue propensity map, the form in which most cryptic-site predictors are evaluated (Section 3.4). The docking outputs exist because a predicted pocket that cannot be handed to the next tool in a workflow is of limited use.

**Figure 2:**
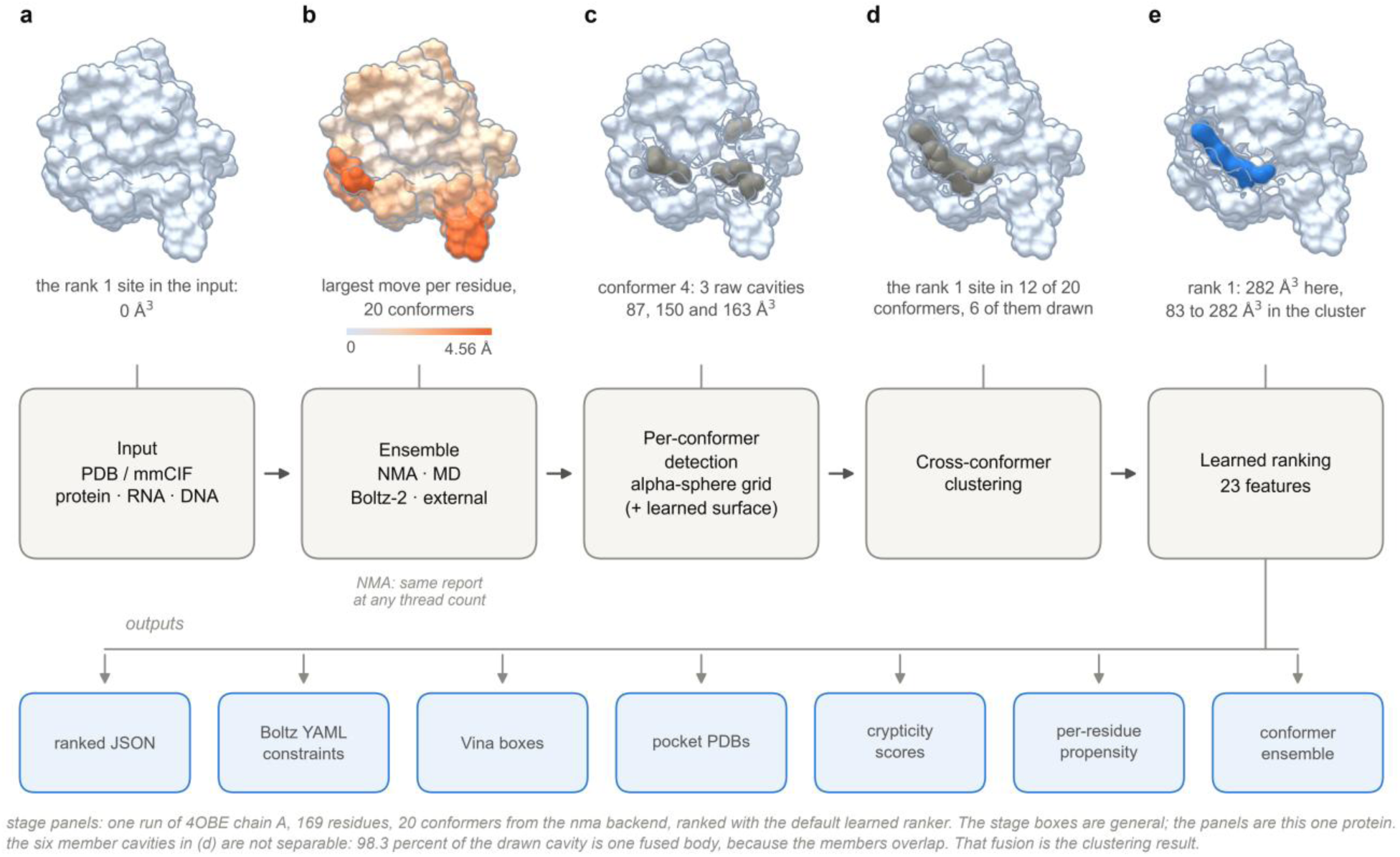
Lacuna’s pipeline, with one run of it above the stages. The schematic is the pipeline: ensemble generation is a plugin point, detection runs independently per conformer, and clustering is what converts transient per-conformer cavities into persistent sites with ensemble statistics. The strip above it is one run of those stages, on chain A of apo K-Ras (PDB 4OBE, 169 residues) with the NMA backend, 20 conformers and the default learned ranker, every panel on one camera at one scale. (a) The input structure. The rank-1 site measures 0 Å³ here, so nothing is drawn for it. (b) The same structure with each residue coloured by how far that residue is ever carried across the 20 generated conformers, 0 to 4.56 Å. This is a displacement field and not an opening field: a residue that moves is not a cavity, and each value is a maximum over the 20 conformers rather than what any single conformer does. (c) One conformer, number 4, with every raw cavity the detector finds in it: three, of 87, 150 and 163 Å³, drawn in a neutral grey because detection has proposed candidates and ranked nothing. (d) The rank-1 site as a cluster, its member cavities drawn from six of the twelve conformers that contain it. The surface is conformer 4’s and five of the six cavities belong to other conformers; the pipeline applies no superposition, so all six are already in that one frame. (e) The ranked site alone, 282 Å³ in conformer 15, in blue. In panels (c) to (e) the surface is left undrawn over the atoms standing in front of the cavities that panel draws, 30.2% of atoms in (c), where all three cavities are drawn, 5.0% in (d) and 4.0% in (e), which is a drawing decision and not an opening in the molecule, and nothing is drawn anywhere the detector found no cavity. The strip is one protein, one run and the default ranker, so it is not evidence for the nucleic-acid path or for the optional rankers; those are described under Design (Section 2.1).

### 2.2 Interface

The command line covers the common case:

~~~
   pip install lacuna-pockets
   lacuna discover protein.pdb
   lacuna discover riboswitch.cif
   lacuna discover protein.pdb --detector surface-fusion
   lacuna discover protein.pdb --backend boltz --conformers 30
   lacuna discover protein.pdb --emit-boltz-constraints --emit-vina-boxes
~~~

Inputs are protein, RNA or DNA structures in PDB or mmCIF format and may come from the PDB, AlphaFold [14], Boltz or Chai. A --homodimer flag builds the biological assembly from BIOMT or _pdbx_struct_oper_list records, which is required for sites at dimer interfaces. A second command, lacuna dock-prep, regenerates docking inputs from an existing pocket report, so a run does not have to be repeated to produce them. Warnings go to standard error, so --quiet leaves standard output clean for scripting, and malformed input is reported in one line rather than a traceback. The Python API exposes the same stages individually for users who want to substitute a component.

Lacuna is also usable without writing a command. A local MCP server exposes two tools to AI assistants such as Claude: one finds cryptic pockets in a local structure file or a four-character PDB identifier, and one exports a docking region as Vina or GNINA boxes or Boltz constraints. The assistant launches it as a subprocess and it runs Lacuna on the user’s own machine, with no hosting, API key or account. The repository also imports as a Claude skill, and hosted servers run Lacuna on Tamarind Bio, Neurosnap and Horus for users who prefer to install nothing.

Lacuna requires Python 3.10 or later. Optional extras (lacuna[openmm], lacuna[boltz], lacuna[plm], or lacuna[all]) gate the heavier dependencies, so a base install pulls only NumPy, SciPy, BioPython and two command-line libraries, and the default pipeline runs with no GPU and no compiled extensions.

## 3 Validation

Unless stated otherwise, every figure in this section was measured with Lacuna 1.5.0, the NMA backend, 20 conformers and the default learned ranker. Results for the optional learned-plm ranker and surface-fusion detector, which need an extra dependency, were measured with version 1.4.0 on their own cohorts of 180 and 184 structures and are marked where they appear; the nucleic-acid results of Section 3.5 were measured with version 1.3.0, except the 1.5.0 sampling cost noted there, and the riboswitch of Section 3.2 was measured with 1.5.0 like the rest of that figure. Confidence intervals are 95% intervals on the paired difference, computed over the structures both arms of the comparison scored.

### 3.1 A worked example

Figure 1 shows Lacuna applied to apo K-Ras (PDB 4OBE, chain A) at default settings. The rank-1 cluster reaches a Jaccard overlap of 0.38 against this target’s benchmark annotation and recovers 79% of its annotated residues, in 0.82 second. The cluster is the emptied nucleotide site rather than the switch-II pocket, and the annotation it is scored against is the P-loop plus switch I, so this example demonstrates the ensemble mechanism rather than recovery of the canonical cryptic site. The switch-II pocket itself is returned at rank 2 and is already open in the input.

The mechanism is visible in the figure rather than asserted. The rank-1 cluster has zero volume in the input crystal structure and reaches 282 Å³ at its widest in the ensemble, and it is detected in 12 of 20 generated conformers. A detector restricted to column 0 has nothing to find. A single target illustrates the mechanism; the aggregate behaviour is in the next section.

### 3.2 A gallery of six targets

Figure 3 puts six targets beside each other, each as its input structure and the conformer in which the site reaches its largest volume. All six ran at the defaults this section uses: 20 conformers, the geometric detector and the learned ranker. Five are proteins sampled with the protein elastic network; the sixth is a 33-nucleotide riboswitch aptamer sampled with the three-bead nucleic network, and nothing else about its run differs, which is what sharing one pipeline means in practice.

**Figure 3:**
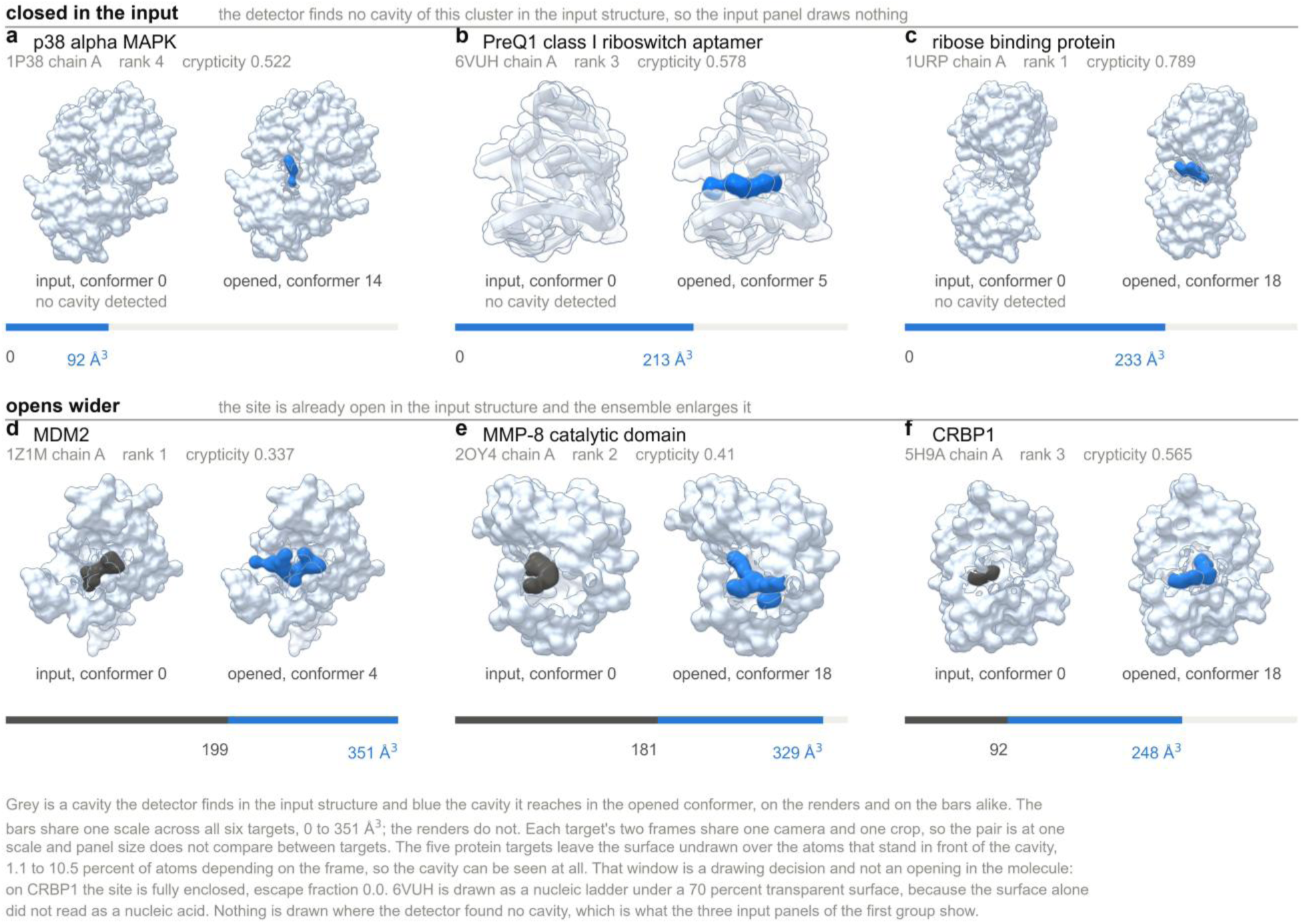
Six targets, input frame beside opened frame. Each pair is one version 1.5.0 run at 20 conformers with the learned ranker: left, the input structure; right, the conformer in which the site reaches its largest volume. Sampling is by normal modes throughout, on the protein elastic network for the five proteins and on the three-bead nucleic network for the riboswitch, which is the network each input calls for. Under each pair, grey is the volume the detector finds at that site in the input structure and blue the volume it reaches in the opened conformer, and the cavities in the renders carry those same two inks. The bars share one scale across all six targets, 0 to 351 Å³; the renders do not, each pair having its own camera and crop. (a) to (c) are the targets with no cavity at the site in the input; (d) to (f) are those where the site is open in the input and opens wider across the ensemble. On MDM2 (d) the opened frame is the widest of a spread: the site measures 199 Å³ in the input and ranges from 90 to 351 Å³ across the ensemble, with a mean of 186.4 Å³, so what the ensemble adds there is the width of its widest conformers rather than a shift of the whole distribution. Each pair of the five protein targets takes one window, the footprint of the cavity in its opened frame, and both of its frames are drawn under that same window, so the two differ only in the molecule; it leaves the molecular surface undrawn over the atoms that stand in front of it, 1.1% to 10.5% of atoms depending on the frame, so that the cavity can be seen at all. That window is a drawing decision and not an opening in the molecule: on CRBP1 (f) none of the 4,000 rays cast from the cavity centroid reaches bulk solvent at the 1.4 Å probe radius the detector itself uses, an escape fraction of 0.0 in both frames drawn. The riboswitch (b) is drawn as a nucleic ladder under a transparent surface, because a surface alone does not read as a nucleic acid at panel size. Nothing is drawn anywhere the detector found no cavity, which is what the three input panels of the first group show.

The six fall into two groups, and both are on claim: a cryptic site is one that is absent or too small to detect in the unbound structure. In the first group the detector finds no cavity at the site in the input structure at all. In the second it finds one and the ensemble opens it considerably wider.

**Table 1.** The six targets of Figure 3.

| Target | Input | Rank | Input volume (Å<sup>3</sup>) | Opened volume (Å<sup>3</sup>) | Crypticity |
|---|---|---|---|---|---|
| **No cavity in the input:** |  |  |  |  |  |
| p38 alpha MAPK | 1P38 A | 4 | 0 | 92 | 0.522 |
| PreQ1 class I riboswitch aptamer | 6VUH A | 3 | 0 | 213 | 0.578 |
| ribose binding protein | 1URP A | 1 | 0 | 233 | 0.789 |
| **Open in the input, wider in the ensemble:** |  |  |  |  |  |
| MDM2 | 1Z1M A | 1 | 199 | 351 | 0.337 |
| MMP-8 catalytic domain | 2OY4 A | 2 | 181 | 329 | 0.41 |
| CRBP1 | 5H9A A | 3 | 92 | 248 | 0.565 |

Source: Version 1.5.0, 20 conformers, learned ranker, normal-mode sampling throughout: the protein elastic network for the five proteins and the three-bead nucleic network for the riboswitch, which is the network each input calls for.
Note: Rank is the site's position in that run's ranked report. Input volume is what the detector measures at the site in the input structure and opened volume what it measures in the widest conformer of the ensemble; crypticity is the score defined under Design (Section 2.1).

Two measurements make the grouping mean something. First, the site of each target was identified rather than assumed: every cluster in the report was scored against that target’s annotated ligand site on centroid distance, residue Jaccard and residue recall, and the cluster shown is the one those measures pick out, which is rank 1 for two of the six and ranks 2 to 4 for the rest. On four of the six it wins all three measures. On the other two it wins two of the three and is nearest the site on both: the cluster shown for p38 has the highest Jaccard but not the highest recall, and the one shown for the ribose binding protein contains every annotated residue but has a lower Jaccard than a smaller cluster covering part of the same cleft. Second, a separate run on the input structure alone, with no conformers generated, confirms what is and is not there, because a cluster’s input volume is a property of the cluster rather than an independent look at the input. For the first group that run finds no cavity at the site; for the second it finds the site open and ranks it near the top.

How often the site is open differs widely across the six, and the report says so per site: persistence runs from 0.048 on p38 and the ribose binding protein, where the cavity exists in a single frame of twenty-one, to 0.952 on MDM2, where it is present in almost all of them. A single-frame cavity is a fact about the measurement and is what the crypticity score is built to express.

The six are what survived a strict bar. Nineteen targets were screened at these settings, and most were set aside on measurement rather than on judgement: the annotated site was already open in the input, so there was no opening to show; or the deposited apo file was not ligand free at the site, so any cavity there is an emptied ligand site rather than a cryptic one; or an annotation was out of register between species; or the run did not recover the site. Each screened target keeps its own measurement directory under paper/figures/software_v3/renders/data, with the commands, the input and report hashes, and the verdict that was reached.

### 3.3 Recovery across four datasets

A site counts as recovered when, among the top five ranked clusters, one reaches a Jaccard overlap of at least 0.25 with the known ligand-contact residues, or one has its centre within 4 Å of the site centroid. The two terms are taken over the top five independently, so a different cluster may satisfy each.

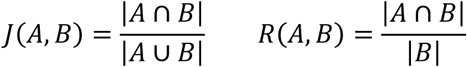

A is the residue set of a candidate’s lining and B the known ligand-contact residues of the target site, so J is the intersection over union that the headline uses and R is plain recall, whose denominator is B alone. Jaccard is used rather than plain recall because recall is size-gameable: a sufficiently large pocket engulfs most of a small known site without being localised on it. CryptoBench figures apply the Jaccard term alone, so that every method in the paired comparisons is scored identically; with the centroid term the same run gives 61.3% (136/222).

All 222 structures of the designated test fold are scored, with nothing skipped. The harness lists every structure it attempts, names any it could not score and counts it as a miss, so the whole fold is the denominator of every figure measured with 1.5.0 here unless that figure states another cohort. Figures carried over from earlier measurements keep the cohort they were measured on and are labelled with it.

The size of that effect determines whether the criterion is doing any work. Ordering the same candidate set by pocket volume alone, with no learned model and no other feature, recovers 74.9% of test-fold structures under a recall threshold of 0.30 but only 53.1% under Jaccard at 0.25, while the learned ranker scores 73.7% under recall. Those figures are on the 180-structure cohort and apply the residue term alone, with the centroid term excluded from each. Measured by recall the trained model cannot be told apart from a volume sort, so every number below uses Jaccard instead. These figures come from benchmarks/verify_recall_gaming.py and are quoted from docs/BENCHMARKS.md, which keeps them as the record of this choice of criterion rather than as a per-release measurement.

#### Cohort note

Several denominators appear in this report, and every figure says which one it carries. The default learned ranker is measured on all 222 CryptoBench test-fold structures. Figures that predate that full-fold measurement stay on the 180-structure cohort the earlier harness scored, which is the single-chain test-fold structures of at most 700 residues: the PLM-assisted ranker, the paired head-to-head comparisons against fpocket and P2Rank, and the criterion-choice figures above. The surface-fusion numbers are a separate five-conformer evaluation on 184 structures, and the per-residue propensity of Section 3.4 is scored over 231 per-residue targets. The companion detector-comparison study [10] uses the 178 on which all four methods it compares produced output, so that every comparison there is paired. The PLM-assisted ranker therefore scores 66.1% (119/180) in this report and 66.3% on that paired 178-structure cohort. The two are the same configuration measured on slightly different sets, not a change in the software. Figures on different cohorts are not compared with each other.

Figure 4 reports recovery on four datasets (Figure 4a). On the designated CryptoBench test fold [15], the largest and most diverse cryptic-site benchmark, the default ranker recovers 59.9% (133/222) and the optional PLM-assisted ranker 66.1% (119/180, measured with version 1.4.0 on the 180-structure cohort). Independent validation is consistent: 69% (31/45) on the PocketMiner set [7], where the PLM-assisted ranker reaches 80% (36/45), and 41% (9/22) on a curated set of apo/holo pairs assembled from the cryptic-pocket literature [2], where both rankers score the same. The split follows CryptoBench’s own homology-separated folds, and the ranker’s coefficients were fitted on the training folds only; no test-fold example entered that fit. The tool as a whole has been developed over many iterations during which test-fold performance was measured, so these numbers are a designated held-out result rather than a claim of full blindness, and the companion study [10] makes the same distinction for the same reason.

**Figure 4:**
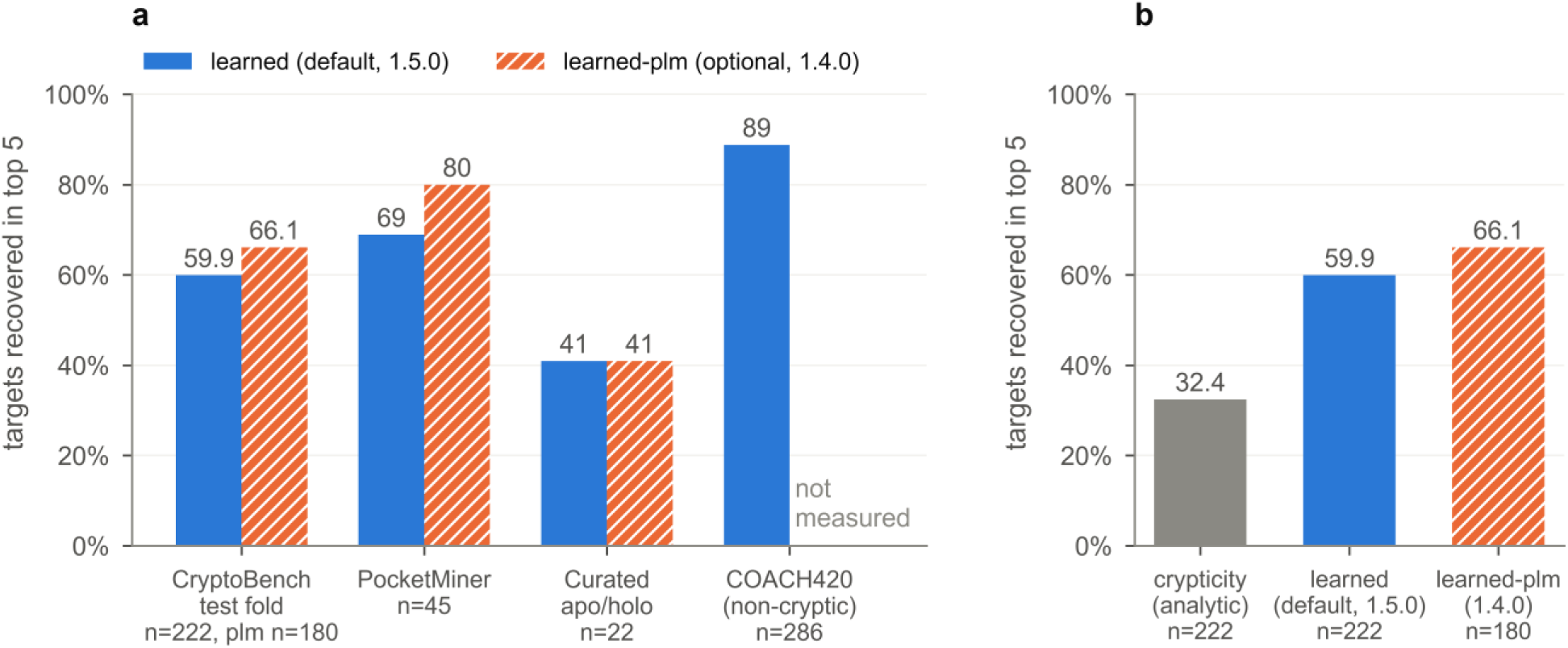
Recovery in the top five predictions, and what ordering contributes. A site counts as recovered at Jaccard ≥ 0.25 against known ligand-contact residues, or a centroid within 4 Å; the CryptoBench bars use the Jaccard term alone. (a) Four datasets. learned is the default ranker, needs only a base install and was measured with version 1.5.0 on all 222 test-fold structures; learned-plm (hatched) additionally requires a protein language model, was measured with version 1.4.0 on the 180-structure cohort, and was not measured on COACH420. CryptoBench values are given to one decimal and the others to whole percent. COACH420 holds general, already-open binding sites rather than cryptic ones and is a control, not a headline: the task is easier and a general-purpose detector is nominally ahead on it. (b) Ranking-strategy ablation on the CryptoBench test fold (n=222). The analytic crypticity rule that served as the pre-1.0 default and the fitted linear model rank the same 1.5.0 candidate set, so only the ordering differs: 32.4% against 59.9%. The PLM-assisted variant (66.1%, hatched) was measured with version 1.4.0 on that release’s candidates and on the 180-structure cohort.

How much of that comes from ordering rather than detection is separable, because the ranking strategy can be changed without touching the pipeline that produces the candidates. Figure 4b varies the ordering, holding the candidate set fixed across the two 1.5.0 bars, which was checked structure by structure over all 222. The analytic crypticity rule that was the default before version 1.0 recovers 32.4% (72/222) of the test fold; the fitted linear model recovers 59.9% (133/222) from the same candidates, a gain of 27.5 points (95% CI +21.2 to +33.8), roughly twice as many, and the PLM-assisted variant 66.1% (119/180, measured with version 1.4.0 on that release’s candidates and on the 180-structure cohort). The detector was already proposing the right pocket for many of the targets the analytic rule missed, and was burying it. This is the single largest effect measured in this report, and it required no change to sampling or detection.

The results above use Lacuna’s default geometric detector, which by construction cannot propose a site that never forms a concavity in any sampled conformer. An optional learned surface detector addresses that class by scoring the probe-accessible surface directly. Pooling the two detectors and ranking the union with a matching fitted model, invoked as --detector surface-fusion, raises top-five recovery from 57.1%, the default’s figure on that set, to 73.9%, and the fraction of structures for which some candidate clears the recovery criterion from 68.5% to 86.4% (measured with version 1.4.0, in a separate held-out evaluation at five conformers on 184 structures, so not directly comparable to the 20-conformer, 222-structure figures above). Within that evaluation the paired gain is +16.8 points of top-five recovery (95% CI +11.4 to +22.8). The surface detector needs the same optional protein-language-model dependency as the PLM ranker and is not the default, so the zero-dependency command is unchanged. This is consistent with the companion study’s finding that no single detector sees all cryptic sites and that their union is larger than any one alone [10].

The fourth dataset is a deliberate control. COACH420 contains holo structures whose pocket is already open, which is an easier task and not the one Lacuna is built for. Lacuna scores 89% (254/286) there with the default ranker. Paired on the 144 structures both tools scored, it reaches 89.6% against P2Rank’s 93.8%, 4.2 points behind with an interval that includes zero (95% CI −9.0 to +0.7). On the same 144 structures the learned ranker is 22.9 points ahead of the analytic druggability rule (CI +15.3 to +30.6), so it is the right default for always-open sites as well as cryptic ones.

The picture on cryptic sites is the same. Every comparison in this paragraph is paired on the 180 CryptoBench test-fold structures the baselines were collected on, so the full-fold 59.9% above is not compared with them. On that cohort the default recovers 57.8% against P2Rank’s 63.3%, 5.6 points behind with an interval that includes zero (CI −12.2 to +1.1), and the PLM-assisted ranker is nominally 2.8 points ahead (CI −4.4 to +9.4; measured with version 1.4.0). No difference between the two tools is resolved at these sample sizes. Against fpocket, the single-structure detector whose lineage Lacuna’s own detection stage follows, the 1.5.0 default is ahead by +14.4 points on those same 180 structures, 104 against 78, with an interval that excludes zero (CI +6.7 to +22.2). The tools also remain complementary rather than interchangeable: the union of the 1.5.0 default with P2Rank recovers 71.7% (129/180) there, with 15 structures recovered by Lacuna alone, and on COACH420 the union of the two reaches 95.8% (138/144). The higher absolute numbers on COACH420 reflect the easier task rather than better performance, and cross-dataset comparison of these figures is not meaningful.

### 3.4 Per-residue propensity

Top-five pocket recovery is what a user reads, but it is not the scale on which this subfield reports: CryptoBench’s own paper, PocketMiner, UniPocket and zPocket all score per residue. The pocket report therefore carries a per-residue cryptic-site propensity:

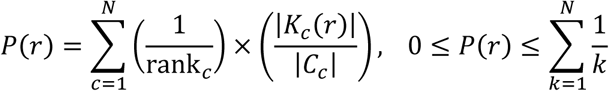

The sum runs over every ranked site in the report, N of them, C_c is the set of ensemble conformers in which site c is detected, and K_c(r) those members of C_c in which residue r lines it; each term is weighted by inverse rank, and the sum is not normalised, so the scale is the harmonic number of the site count rather than a probability. The propensity is measured here on the CryptoBench held-out test fold, over its 231 per-residue targets and 76,633 residues, which is a different cohort from the 222 structures the top-five tables above use, with labels taken as the union of the per-chain apo pocket selections, and the aggregation form was chosen on the training folds and then applied unchanged (Table 2).

**Table 2.** Per-residue cryptic-site propensity on the CryptoBench test fold.

| Scorer | AUROC | AUPRC |
|---|---|---|
| Propensity, 20 conformers | **0.766** [0.737, 0.793] | **0.340** [0.308, 0.373] |
| Propensity, input structure only | 0.697 | 0.249 |
| Burial neighbour count | 0.586 | 0.115 |
| Random expectation | 0.500 | 0.067 |

Source: Version 1.5.0 on the CryptoBench held-out test fold, 231 targets and 76,633 residues, except the burial row, which does not use the ensemble and was measured with 1.4.0.
Note: Precision is averaged within each protein and then over proteins, so large proteins do not dominate; brackets are 95% confidence intervals over targets.

Two rows carry the argument. The first is the input structure: the same scorer on the deposited coordinates alone reaches AUPRC 0.249 against 0.340 from the ensemble, a paired difference of +0.092 (95% CI +0.069 to +0.115), and the gap survives a stricter label that counts only the main holo partner, 0.304 against 0.216. This is the most direct evidence in this report that the ensemble carries information the input structure does not. The second is burial: a score built from cavity lining could be a restatement of how buried each residue is, which would make it a geometric re-description rather than new information. Paired over targets, the propensity beats the burial control by +0.180 AUROC and +0.225 AUPRC.

Published comparators need the averaging convention stated with them. CryptoBench’s table pools every residue into one curve, and pooled the same way this scorer gives AUROC 0.807 and AUPRC 0.227: level with P2Rank on AUROC and above it on AUPRC (0.81 and 0.21), above PocketMiner on both (0.76 and 0.19) on their reported subsets, and below CryptoBench’s own sequence model (0.86 and 0.36), which is the expected place for a geometric, ensemble-derived score carrying no learned sequence component. The subsets differ: CryptoBench’s figure is the whole test set, PocketMiner’s drops the structures it could not process, and P2Rank’s is the single-chain apo subset.

An independent evaluation prompted this output. zPocket [18], winner of the Octavian-Eugen Ganea Prize for Best Paper at MoML 2026, benchmarked Lacuna on their PocketWatch dataset and placed the surface-fusion variant’s detection ceiling highest of any single detector they tested, 0.780, with the default detector at 0.616, in an evaluation of a build predating version 1.4.0. That reproduced independently the coverage-versus-conversion pattern reported in Section 3.3 and in the companion study [10]. The ceilings are not like-for-like, because the two tools emit very different numbers of candidates per protein, 57.5 to 64.9 centroids against 1.1, with Lacuna’s report cap raised to 1,000 for that benchmark.

The scope is narrow and worth stating: this measures where cryptic sites are, not how well candidate pockets are ordered, and it leaves the ranker untouched.

### 3.5 RNA and DNA

Nucleic-acid input needs no extra flags. The input’s residues select the three-bead elastic network described under Design (Section 2.1), and detection, clustering, ranking and output are the same code the protein path runs. The one difference is in the ranker’s weight vector: on input with no protein in it a single duplicated burial weight is zeroed, because its fitted sign is correct for protein, where the ligand site is the shallower candidate, and backwards for nucleic acid, where it is the more buried one. The other 22 weights are shared and nothing is refitted. The recovery figures below are measured on RNA; the DNA path runs that same code and is exercised for reproducibility (Section 3.6), and no DNA ligand-site recovery benchmark is reported here.

Across 31 RNA structures drawn from 12 independent BGSU equivalence classes and spanning 62 to 459 nucleotides, a single pass finds the ligand site for 87% of structures. The case the ensemble exists for is the subset with no detectable cavity in the input at all: there the ensemble takes the fraction of sites found from 44% to 73%, measured across 62 apo structures from 41 sequence families. Both figures were measured with version 1.3.0.

Ranking on short RNA is reported against a random-ordering baseline in the repository’s documentation rather than as a bare top-five percentage, because on a structure with only a handful of candidates a top-five hit is close to free. Since 1.5.0 nucleic-acid sampling runs through the same loop as protein and at the same cost: on a 1,483-bead protein and nucleic-acid complex, ensemble generation for 20 conformers takes 6.4 seconds and peaks at 671 MB (Section 3.7).

### 3.6 Reproducibility

Since version 1.5.0 the default NMA backend writes an identical pocket_report.json for the same input and options at any BLAS thread count, and between a Windows and a Linux build on the 40 proteins compared. There is nothing to configure: no thread variable to pin and no seed to pass.

The report was compared byte for byte between the default thread count and a single thread on 30 inputs, spanning 6 proteins of 122 to 3,724 residues, 17 RNA structures, 2 DNA structures, 2 protein and nucleic-acid complexes, 2 symmetric assemblies built with --homodimer, and the K-Ras example of Figure 1: 30 of 30 identical, with the conformer coordinates behind those reports agreeing to float32 rounding. Across 184 structures at 1, 2, 8 and the default number of threads, and between the Windows and the Linux build, at most 40 of 26 million coordinates differed, none by more than 1.9e-6 Å, which the report’s rounding removes. The K-Ras run behind Figure 1 reproduces on the same terms, byte-identical over 13 runs at the default thread count and pinned to one thread.

Two boundaries belong with the claim. The guarantee is for the nma backend; the openmm and boltz backends depend on their own libraries and hardware. And one class of input falls outside it, which Lacuna detects and reports: if the elastic network is under-constrained, so that it has no vibration to sample, the run warns and still writes a report, but that ensemble is not physically meaningful and is not reproducible between machines. In practice this is a predicted model whose disordered regions are drawn as long extended ribbons.

Results are comparable within a release rather than across releases: reports differ from those version 1.4.0 wrote for the same input, because every ensemble is drawn by the current sampler and a site’s lining follows the majority rule described under Design (Section 2.1). Earlier structures should be re-run with 1.5.0 before being compared with new ones.

### 3.7 Runtime and memory

All timings in this section were measured on one desktop machine running Windows 11, an AMD Ryzen 9 7900X with 12 cores, and no GPU.

On 30 single protein chains of 57 to 676 residues, Lacuna 1.5.0 runs the full pipeline, ensemble generation through ranking, in a median of 1.9 seconds per chain with the NMA backend at 20 conformers, with linear algebra pinned to one thread (Figure 5). The chains are single extracted chains drawn across the CryptoBench size range rather than cherry-picked. The fastest chain takes 0.4 second and the longest, at 676 residues, 5.5 seconds; each point is the median of three runs. A power law fitted to the 30 points gives an exponent of 1.05, so cost grows close to linearly with chain length.

**Figure 5:**
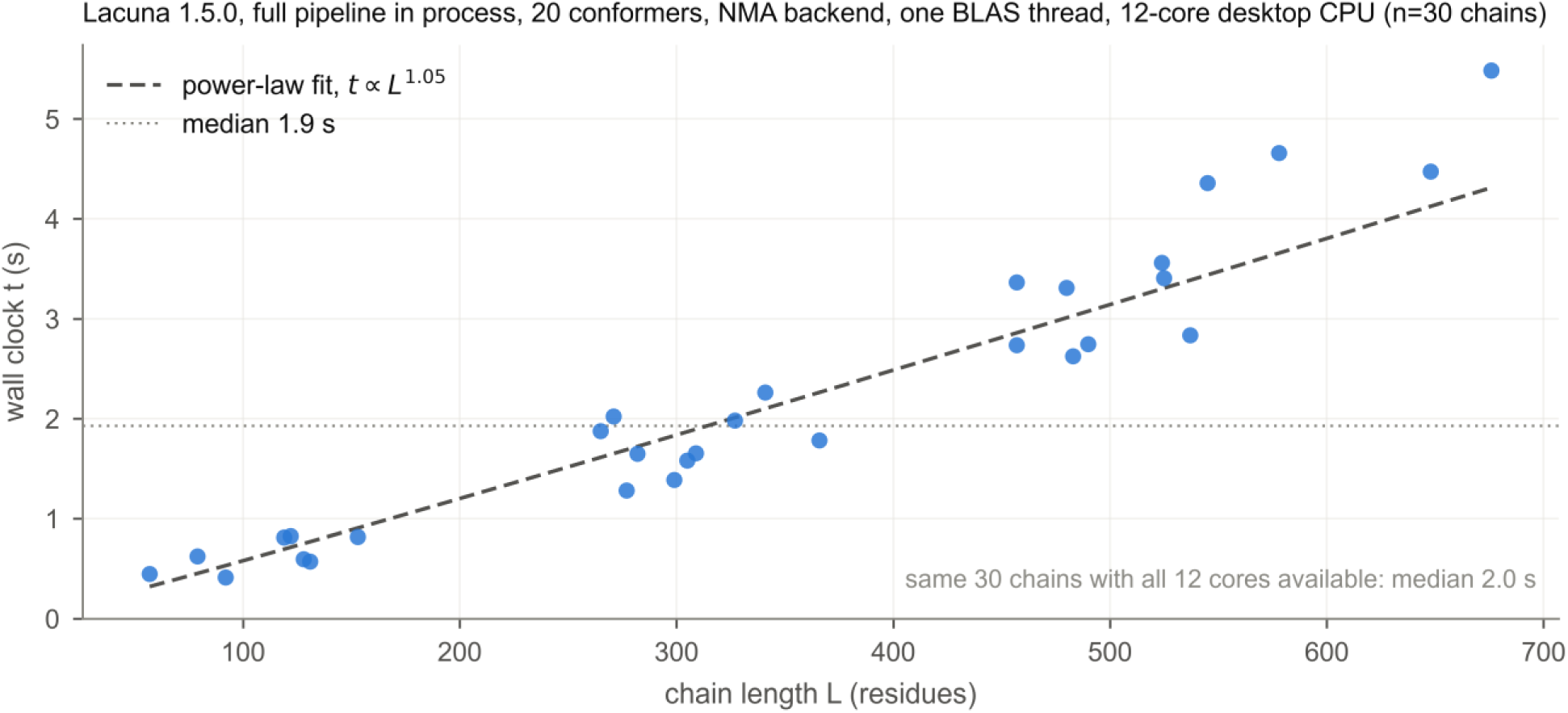
Wall clock against chain length. Lacuna 1.5.0 running the full pipeline, ensemble generation through ranking, at 20 conformers with the NMA backend, on 30 single protein chains of 57 to 676 residues. One BLAS thread, 12-core AMD Ryzen 9 7900X desktop running Windows 11, no GPU. Each point is the median of three runs, timed in one warmed interpreter, so CPython start-up and package imports are outside the measurement. Dashed line, power law fitted to the 30 points, exponent 1.05. Dotted line, median 1.9 seconds. With all twelve cores available the same chains give a median of 2.0 seconds, and the same number of pockets for all 30.

Timings are of the pipeline itself, measured in one warmed interpreter. Starting a fresh Python interpreter and importing Lacuna’s dependencies adds about 0.75 second to a single command-line invocation, which a user pays once per run and a script once per session. With all twelve cores available the same 30 chains give a median of 2.0 seconds, and the per-chain difference between one thread and twelve has a median of 0.01 second, so nothing is lost by running single-threaded; with the NMA backend the pocket report is identical at either thread count.

On the reference proteins of the repository’s speed table the same pipeline takes 0.59 second at 122 residues, 0.84 second for K-Ras chain A at 169 (0.82 second pinned to one thread, the run behind Figure 1), and 5.91 seconds at 536, measured at 20 conformers on a 12-core desktop CPU as the median of three runs, and is 7% to 30% faster than 1.4.0 across that table.

Ensemble generation dominates memory. At 20 conformers the 1.5.0 sampler peaks at 105 MB for 339 residues and 1,202 MB for 3,724, against 154 MB and 3,563 MB for the same structures with 1.4.0 on the same machine, and generation time falls from 0.89 to 0.23 second and from 73.4 to 49.5 seconds across that range (Figure 6). From roughly 1,300 residues up the sampler needs less than half the memory of the previous release and about a third at 3,700, which is what brings assembly-sized inputs inside an ordinary desktop. RNA and DNA are sampled through the same loop: on a 1,483-bead protein and nucleic-acid complex, peak memory for 20 conformers is 671 MB and generation takes 6.4 seconds. These figures are for ensemble generation alone, not for the whole pipeline.

**Figure 6:**
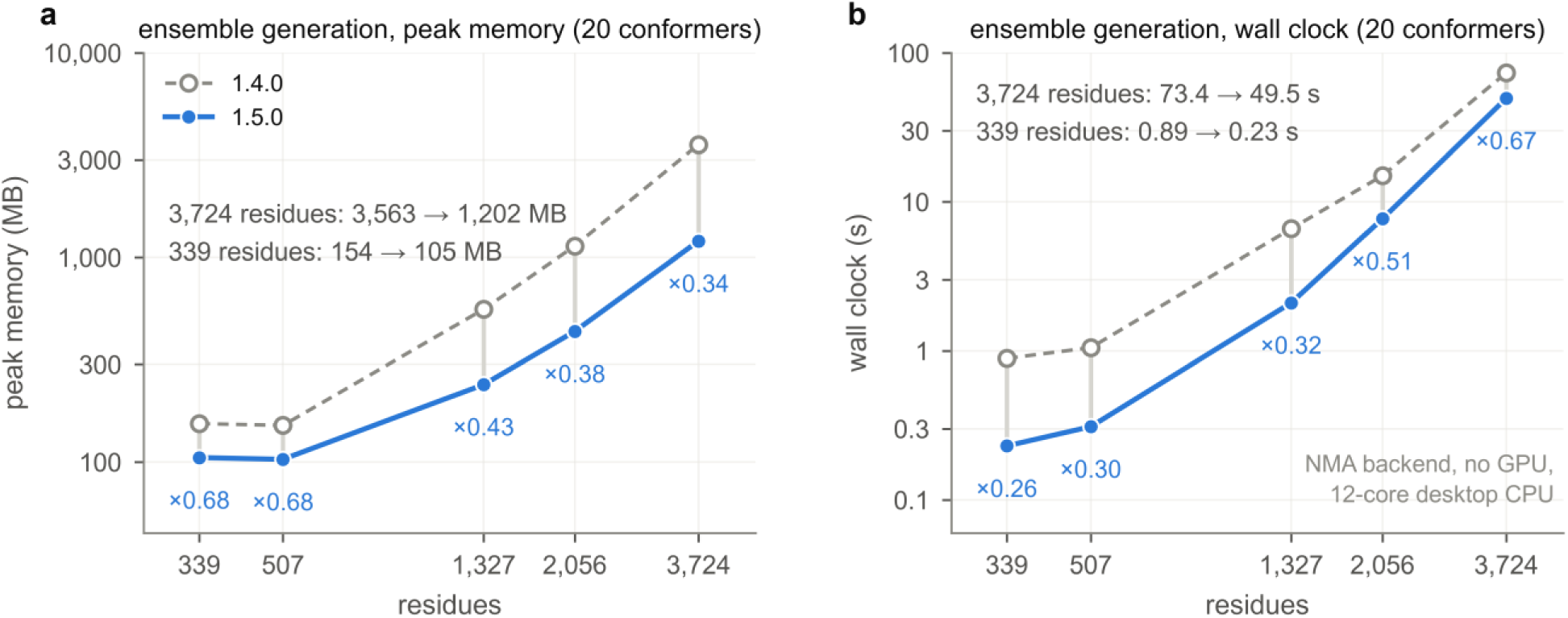
Ensemble generation at 20 conformers with the NMA backend: peak process memory (a) and wall clock (b) for Lacuna 1.4.0 and 1.5.0, timed back to back on the same 12-core desktop CPU with no GPU. Axes are logarithmic. The label under each 1.5.0 point is its ratio to 1.4.0 at the same size. Memory falls to about two thirds at 339 and 507 residues and to about a third at 3,724. Values as published in the release benchmarks for 1.4.0 and 1.5.0.

The practical consequence is that ensemble-based pocket discovery on a single protein is an interactive operation rather than a scheduled job, which is the design goal. The heavier backends are correspondingly more expensive: the molecular dynamics and Boltz-2 backends cost minutes to hours per target and are appropriate when the default’s harmonic sampling is known to be insufficient.

## 4 Conclusions

Lacuna packages ensemble-based cryptic pocket discovery as a tool that installs with pip and runs in seconds, rather than as a protocol requiring a simulation budget. Its distinguishing choices are that the ensemble backend is a swappable parameter, that detections are clustered across conformers into sites carrying ensemble statistics, and that ranking those sites is a fitted problem rather than a hand-designed rule.

It now handles RNA and DNA with no extra flags, gives the same report on every machine, adds a per-residue propensity map that places it on the scale the subfield reports, and is available to AI assistants through a local MCP server and on hosted platforms that need no installation. Measured with 1.5.0 on CryptoBench’s designated test fold, the zero-dependency default recovers 59.9% of all 222 structures of that fold in its top five, and the per-residue propensity reaches AUPRC 0.340 against 0.249 for the same scorer on the input structure alone. On 30 single chains of 57 to 676 residues the full pipeline takes a median of 1.9 seconds per chain on one thread. The optional PLM ranker and the pooled detector reach 66.1% on the 180-structure cohort and 73.9% on the separate 184-structure evaluation, both measured with version 1.4.0 as reported above.

### 4.1 Limitations

#### Two limits are worth stating plainly

The first is ranking. Across the candidate set Lacuna generates, some cluster clears the recovery criterion for 76.6% of CryptoBench test-fold structures (170/222), against the 59.9% the default ranker places in the top five. The site is often found and then out-ranked, and part of the loss sits just outside the cutoff: the correct cluster is at rank 6 to 8 for 13 structures, and top-eight recovery is 65.8% (146/222). The optional surface detector raises both ends, to 86.4% found and 73.9% ranked into the top five, in that separate five-conformer evaluation on 184 structures, but the gap between them persists. Ranking is the largest gap that closes without changing the sampler, and it is not the larger loss: 23.4% of test-fold structures (52/222) carry no qualifying candidate at all, which is the sampling limit stated below. Several attempts to close that gap returned nothing measurable: spatial non-maximum suppression, merging adjacent sub-pockets, hard-negative mining, gradient boosting in place of the linear model, and importing P2Rank’s own per-pocket confidence as a ranking feature. The last is the most informative: if per-point scoring were the missing ingredient, handing the ranker P2Rank’s opinion directly should have helped, and it did not.

The second is sampling. The default elastic network is harmonic and cannot generate large hinge or interface openings, so a site whose opening requires one will not appear in any sampled conformer and cannot be detected or ranked. Enhanced-temperature dynamics, metadynamics along an apo-derived collective variable, and scaled-water dynamics were each null against baseline at single-workstation sampling. Catching those events reliably appears to need tens to hundreds of nanoseconds across dozens of replicas per target, which is a cluster-scale cost rather than a missing algorithm. This is the gap the co-folding backend is intended to probe, since a generative model samples conformational diversity without paying for the trajectory between states; that backend remains experimental.

## Data and code availability

Source code is at https://github.com/mooreneural/lacuna under the MIT licence, released as v1.5.0 and installable as pip install lacuna-pockets. The exact version described here is archived on Zenodo at doi:10.5281/zenodo.24239150 [9]; the concept DOI doi:10.5281/zenodo.20533638 resolves to all archived versions. All benchmark scripts, including those that produce every number in Figure 4, are in the repository’s benchmarks/ directory and download their datasets automatically; the per-residue figures of Table 2 are reported in docs/BENCHMARKS.md. The runtime measurements of Figure 5 are reproduced by the sweep script and provenance notes in paper/figures/software_v3/; Figure 6 plots the sampler table published in docs/BENCHMARKS.md. The rendered panels of Figures 1, 2 and 3 are recorded in paper/figures/software_v3/PROVENANCE_renders.md, which gives for each panel the command that produced the measurement behind it, the input and report hashes, the voxel export and its checks, and the renderer settings; the per-target measurement directories it points at carry the rest.

The local MCP server is in mcp_server/ and the Claude skill in skills/lacuna/, both in the same repository. Hosted, no-install deployments run Lacuna on Tamarind Bio (https://app.tamarind.bio/tools/lacuna), Neurosnap (https://neurosnap.ai/service/Lacuna%20Cryptic%20Pocket%20Discovery) and Horus (https://os.templecompute.com/library/lacuna-cryptic-pocket-discovery-2500e2).

